# Social avoidance of mice in pain in naturalistic conditions

**DOI:** 10.1101/2023.10.31.564932

**Authors:** Olivia Le Moëne, Max Larsson

**Author notes:** **Corresponding author**: Olivia Le Moëne **Email:**. **ORCIDs**: OLM (0000-0003-3599-0211), ML (0000-0003-3584-7829).

## Abstract

Pain and social behavior are subject to reciprocal modulation. Both humans and rodents experience emotional contagion from afflicted conspecifics, and may act to relieve the afflicted state of these. Little has been done to investigate the motivation of such prosocial behavior in rodents in naturalistic conditions. Here, we analyzed social interactions in mice group-housed in a seminatural environment (SNE). Social buffering reduced nocifensive behavior in formalin-injected mice. These mice were also both socially withdrawn and avoided by other mice. These findings appear counter to those showing empathy in mouse pain models. It is possible that in naturalistic conditions, healthy mice simply avoid individuals in pain and the cost associated with emotional contagion. Interestingly, behaviors involving direct body contact were not different between saline– and formalin-treated mice, and thus may carry a prosocial, altruistic component. These findings unveil new patterns of social modulation by pain in a naturalistic laboratory setting holding high translational value.

**Teaser:** In a new, ethological assay, mice in pain are socially withdrawn and avoided by others, challenging findings of empathy in rodent models.

## 1. Introduction

Social interaction can have a mitigating effect on pain in humans [1]. Therefore, it has been suggested that such an effect could also be found in non-human animals. Indeed, multiple studies have defined 1) a pain-modulating effect by social behavior and 2) remodeling of social interactions by pain. One of the most well-established effects of social interaction, known as social buffering, is the alleviation of pain behavior by the presence of a healthy conspecific [2, 3], through the down-modulation of the HPA axis stress response [4]. Social buffering is most effective between familiar individuals co-housed for several days [3, 5], making social closeness a key feature in interaction between social behavior and pain. However, most studies have compared unfamiliar mice or cagemates in simplified, reduced social settings, that hold little external validity [6]. More naturalistic environments have been implemented to map social relationships in a more realistic way, whether focusing on affiliative [7], aggressive [8] or sexual behaviors [9]. To our knowledge, pain and social behavior in rodents have not yet been investigated under such naturalistic conditions.

Interestingly, healthy animals often exhibit heightened nociception following exposure to a suffering conspecific [10, 11]. Emotional contagion is suggested to constitute the most basic manifestation of animal empathy and has been demonstrated in a variety of mammals, including laboratory rodents [12]. Though emotional contagion is well established in laboratory rodents, the translation of emotional contagion, or an empathic feeling, in prosocial behaviors toward an afflicted conspecific is less studied. Those behaviors may carry altruistic value, if motivated by the improvement of the other’s internal state.

One of the most commonly observed affiliative behaviors in mammals is allogrooming, a form of social grooming defined as “caregiving through physical contact” [13]. Though this behavior can be directed to any body part on the receiver, it is often performed in the nape of the neck, where it is commonly observed in equids, primates or rodents. This area is believed to be a target of choice as it is one of the least vulnerable to physical attack, limits eye contact between the groomer and the receiver, and is difficult to reach by oneself [14]. Therefore, the neck is a highly socially relevant area to study affiliative touch. Cutaneous rodent pain models, such as the formalin assay, are mostly directed toward the plantar hind paw rather than more socially relevant areas. However, measures of paced social interactions are dependent on animal mobility, which can be affected by limb pain. A new, neck-targeted pain model would be justified to investigate pain and social behavior while avoiding the confound of reduced locomotion and could unveil critical differences from traditional experimental settings.

This study aimed to characterize the interaction between pain and social behavior in mice hosted in naturalistic conditions. We hypothesized that mice in pain would receive increased social attention and allogrooming. Due to the presence of several familiar conspecifics and thus of social buffering, we expected the mice to show attenuated nocifensive behavior.

## 2. Results

### 2.1 Formalin neck injection elicits a typical pain face

In order to validate the nape of the neck as a target for pain induction, we injected mice with either saline (S, 9mg/ml, 2 µl) or formalin (F, 2% in saline, 2 µl) in either the traditional plantar hind paw (H) site or the nape of the neck (N) (Fig.1). We first established the facial expression elicited by treatment in the nape of the neck in mice placed in an individual observation cubicle. During the first phase of formalin, mice showed a pointier snout (p < 0.001) and a more convex face inclination (both parameters reduced, p < 0.001 and p = 0.029, respectively), in combination with the ears being directed backwards and placed on the side of the head (both ear position and ear opening increased, p < 0.001 and p = 0.027, respectively), but no significant effect on eye opening (p = 0.137) (Fig. 2A). During the second phase, the face remained more convex (p = 0.007) and eye opening was significantly smaller (p = 0.005). Other parameters returned to their baseline levels (all ps > 0.123). Response profile for SN showed a slight reduction in eye opening (p = 0.006) and face inclination values (p = 0.009). In summary, this facial response profile was similar to that observed after formalin injection into the hind paw [15]. Interestingly, response profiles to formalin injection tended to be more pronounced for females, as well as in measures of the left side of the face. However, direct statistical comparisons of facial parameters values showed little to no sex nor side difference (Fig. S2).

**Figure 1.**
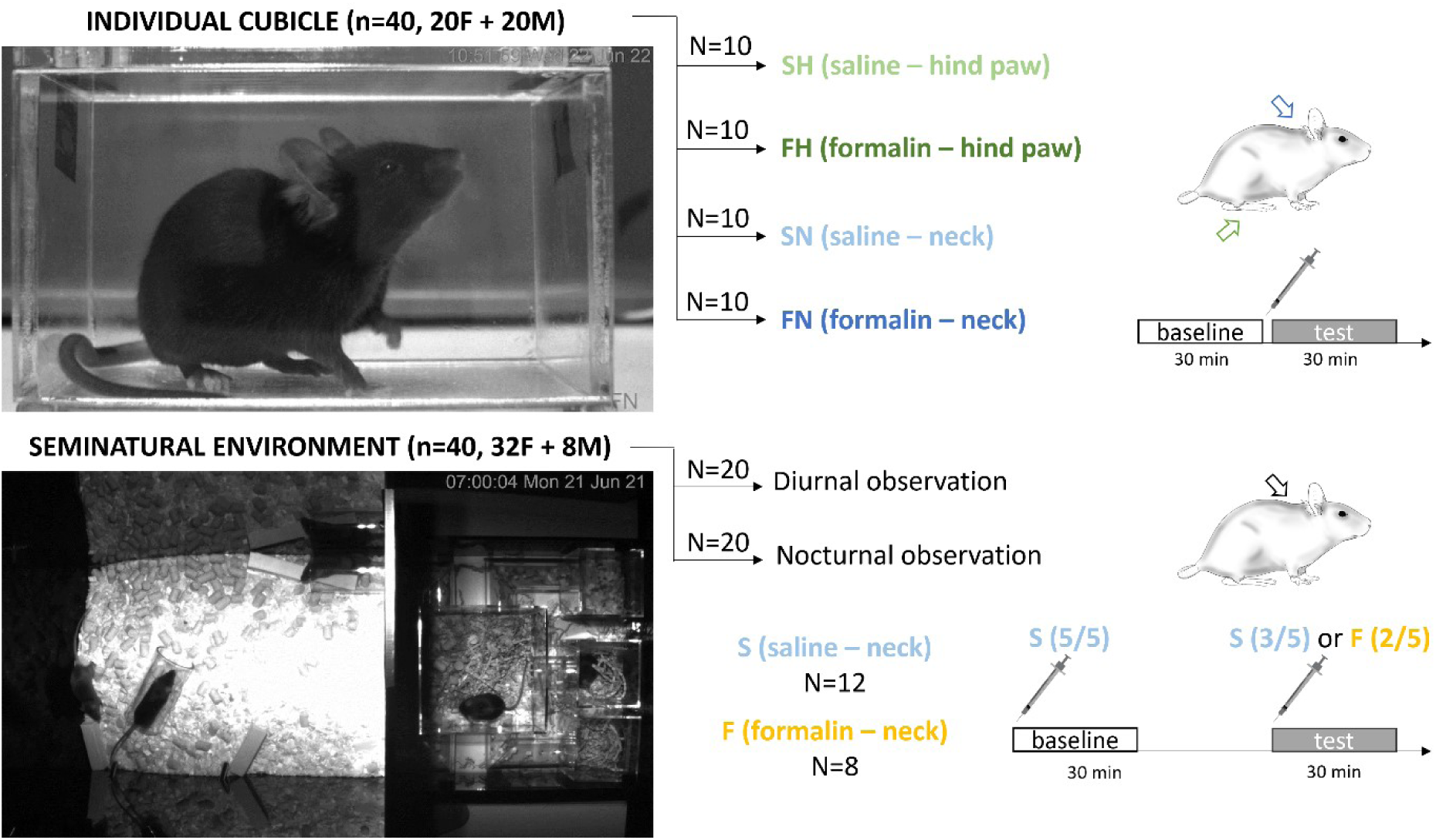
Schematic protocol of the experiment. Individual observation: mice were singly placed in an observation cubicle and after 30 min of baseline injected with either saline (9mg/ml, 2µl) or formalin (2%, 2µl) into either the hind paw or the nape of the neck. Individual observation took place during the diurnal phase of the light cycle. Naturalistic observation: eight groups of five mice (4 females + 1 male) were placed in a seminatural environment. They were observed following saline injection into every individual, and 1 hour later following formalin injection into 2 of them. Four groups were observed during the diurnal phase of the light cycle, and four groups during the nocturnal phase.

**Figure 2.**
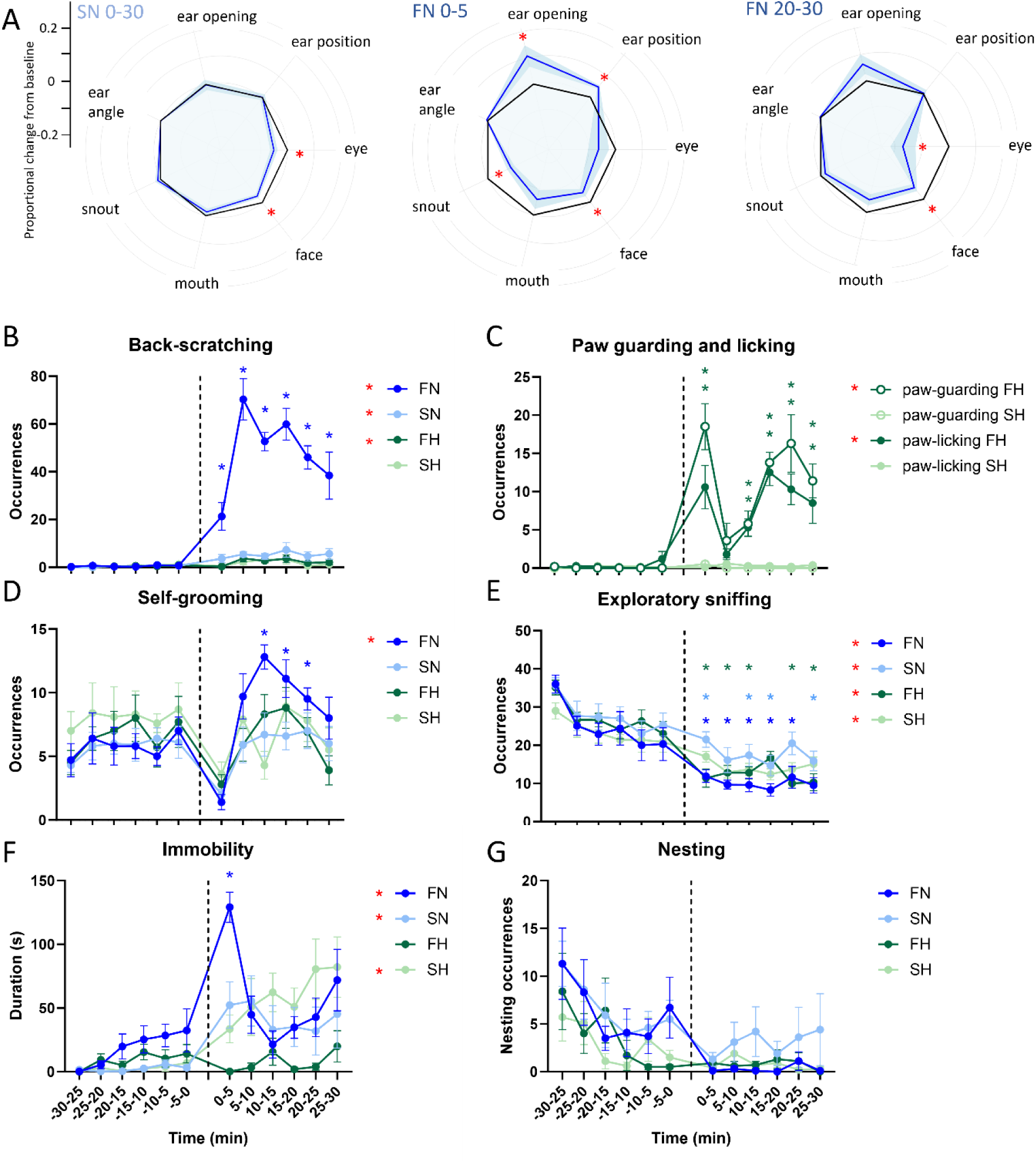
Facial expressions and behaviors surrounding saline (light colors) and formalin injection (dark colors) into the right hind paw (green colors) or the nape of the neck (blue colors) in mice hosted individually. **A**. Response profiles based on proportional change from average baseline value for each facial parameter. Profiles for injection into the neck of saline (30 min average, SN 0-30) and of formalin (first 5 min, FN 0-5; 20 to 30 min post-injection, FN 20-30). Facial parameters are abbreviated as such: eye = eye opening, snout = snout position, mouth = mouth position and face = face inclination. Red stars indicate a significant difference from baseline based on a one-sample *t*-test. **B** Back-scratching. **C**. Paw-guarding and paw-licking. **D**. Self-grooming. **E**. Exploratory sniffing. **F**. Immobility. **G**. Nesting. Two-way ANOVAs for repeated measures. Red stars indicate a main effect of the injection compared to baseline for the adjacent experimental group. Other colored stars refer to a significant difference from the matching baseline time interval, according to Šídák’s post hoc tests. Star color matches that of the experiment group it applies to. SH, n=10; FH, n=10; SN, n=10, FN, n=10. *, p < 0.05. Data are mean ± SEM.

### 2.2 Absence of a two-phase pattern following formalin neck injection

Behaviors in the observation cubicle were monitored over the baseline and following injection for the FN (formalin-neck), SN (saline-neck), FH (formalin-hind paw) and SH (saline-hind paw) groups. For FN, back-scratching was overall largely increased after injection compared to baseline (F_(1.000, 9.000)_ = 87.14, p < 0.001) (Fig 2B). Notably, back-scratching was strongly increased at all time intervals (all ps < 0.031; interaction: F_(3.366, 30.30)_ = 11.55, p < 0.001). It was slightly increased after injection also for SN and FH (ps < 0.016).

Consistently with the effect of FN on back-scratching, self-grooming also increased after injection for the FN group (F_(1.000, 9.000)_ = 8.172, p = 0.019). This increase was significant from 10 to 25 min following formalin neck injection (all ps < 0.036; interaction: F_(2.702, 24.32)_ = 4.743, p = 0.012) (Fig.2D).

Paw-licking and paw-guarding were only observed following injections into the paw. Therefore, only SH and FH groups are analyzed. Both behaviors were increased following formalin, but not saline injection (FH: ps < 0.001, SH: ps > 0.200). Their temporal dynamics showed the typical two-phase pattern: paw-guarding and paw-licking were increased during the first 5 min and then again between 10– and 30-min following formalin injection (ps < 0.050; interactions: ps < 0.008) (Fig.2C).

Upon re-introduction to the cubicle after injection, sniffing was reduced for all groups (all ps < 0.002). We found several interactions as visualized in Fig. 1E. In parallel to sniffing reduction, immobility increased for SN (F_(1.000, 9.000)_ = 7.442, p = 0.023), SH (F_(1.000, 9.000)_ = 32.40, p < 0.001) as well as FN (F _(1.000, 9.000)_ = 28.54, p = 0.001) but not FH (F_(1.000, 9.000)_ = 0.3436, p = 0.572). In particular for FN, immobility was increased in the first 5 min following injection only (p < 0.001; interaction: F_(2.896, 26.06)_ = 9.039, p < 0.001) (Fig.2F). There were few nesting occurrences in the SH and the FH groups, thus this behavior was not analyzed but simply illustrated in Fig.2G.

### 2.3 Nocifensive back-scratching is reduced in the presence of conspecifics

After characterizing facial expressions and nocifensive behaviors following formalin injection into the neck compared to the traditional hind paw, we analyzed the effect of the social environment on these behaviors. Neck injections of saline and formalin were thus given to mice hosted in groups in a seminatural environment (SNE). We compare the data obtained in the cubicle and presented in Fig. 2 with the new data obtained in the SNE. Data from the 30-min baseline was averaged to obtain a single baseline value in both apparati. We observed mouse behavior during the diurnal phase in the SNE, as we did for mice observed in the cubicle.

First, we compared formalin-injected mice in both apparati (FN-cubicle and FN-SNE). In the SNE, the increase in back-scratching was comparatively lower than it was in the cubicle (30.1 ± 3.2 vs. 48.1 ± 3.4 occurrences/5-min respectively, F_(1, 16)_ = 6.27, p = 0.024). Back-scratching in the cubicle was higher than at baseline at all time intervals (ps < 0.035), but only 5 to 10 and 20 to 25 min post-injection in the SNE (p = 0.049 and p = 0.015, respectively; interaction: F_(6, 96)_ = 2.97, p = 0.011) (Fig.3A). Comparatively, self-grooming was also less frequent in the SNE than in the cubicle (F_(1, 16)_ = 6.316, p = 0.023) (Fig.3B).

**Figure 3.**
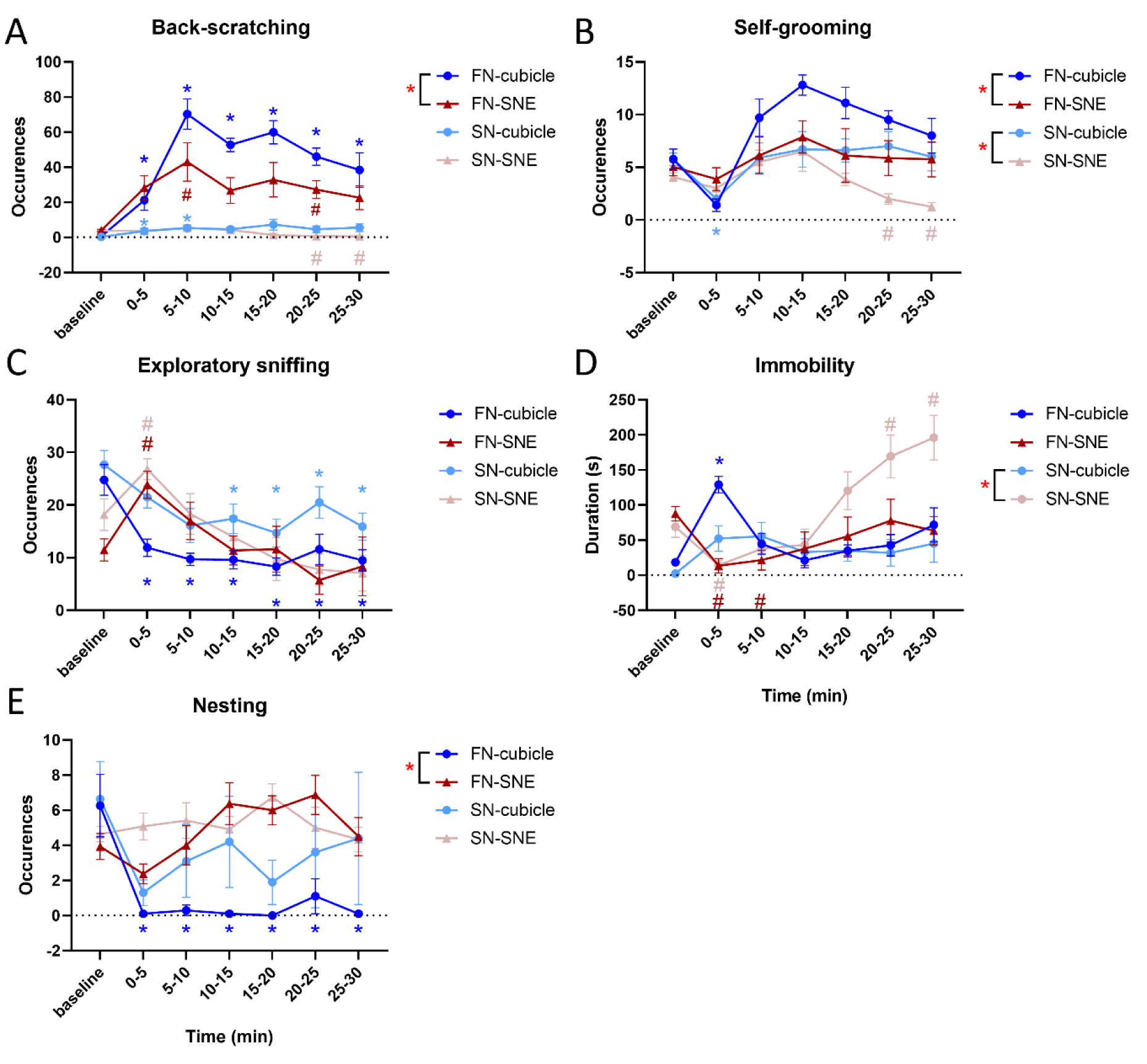
Nocifensive and maintenance behaviors surrounding saline (light colors) and formalin injection (dark colors) into the neck in mice hosted in an individual cubicle (blue colors) or a seminatural environment (SNE, red colors) during the diurnal phase. FN-cubicle and SN-cubicle data are re-plotted from Fig.2 using a single average baseline value. A. Back-scratching. B. Self-grooming. C. Exploratory sniffing. D. Immobility. E. Nesting. Red stars in the legend indicate a main effect of the apparatus (cubicle versus SNE). Two-way ANOVAs for repeated measures on the time interval factor. Red stars indicate a main effect of the apparatus. Symbols on plots indicate significant differences from matching baseline value for animals tested in the cubicle (*) or in the SNE (#) according to Šídák’s post hoc tests. Symbol color matches that of the experimental group it applies to. SN-cubicle, n=10; SN-SNE, n=12; FN-cubicle, n=10; FN-SNE, n=8. Data are mean ± SEM.

Exploratory sniffing frequency was overall similar in the cubicle and the SNE (F_(1, 16)_ = 0.103, p = 0.7526). However, while exploratory sniffing was increased in the first 5 min upon return to the SNE (interaction: F_(6, 96)_ = 4.944, p < 0.001, post-hoc comparison: p = 0.010), it was reduced at all time intervals following re-introduction in the cubicle (ps < 0.005) (Fig.3C). Conversely, immobility was almost abolished in the first 10 min upon return to the SNE (interaction: F_(6, 96)_ = 6.347, p < 0.001, post hoc comparisons: ps < 0.033) while it peaked in the first 5 min upon return to the cubicle (post hoc comparison: p < 0.001), though not being significantly different between the apparati (F_(1, 16)_ = 0.005, p = 0.943) (Fig.3D). Finally, nesting behavior was more frequent in the SNE than in the cubicle (F_(1, 16)_ = 49.76, p < 0.001). In the cubicle, nesting was almost absent at all time intervals post-injection (all ps < 0.001; interaction: F_(6, 96)_ = 6.341, p < 0.001) (Fig. 3E).

We also compared saline-injected mice in the SNE and the cubicle (SN-cubicle and SN-SNE). Back-scratching showed some variation along the time intervals (interaction: F_(6, 120)_ = 2.843, p= 0.013, see Fig.2A) but no main effect of the apparatus (F_(1, 20)_ = 3.203, p= 0.089). As for formalin-injected mice, self-grooming was less frequent in the SNE than in the cubicle for saline-treated mice (F_(1, 20)_ = 10.09, p = 0.005) (Fig.2B). Saline-treated mice showed the same pattern of interaction for exploratory sniffing as formalin-treated ones (interaction: F_(6, 120)_ = 2.513, p = 0.025). Namely, in the SNE, sniffing peaked in the first 5-min upon return, while in the cubicle it decreased after 10 min (Fig.2C). Finally, immobility lasted longer in the SNE (F_(1, 20)_ = 11.080; p = 0.003) and increased overtime (interaction: F_(6, 120)_ = 7.173; p < 0.001). Post hoc comparisons for the reported interactions for each behavior are visualized in Fig.3A-E.

### 2.4 Social avoidance of mice in pain

Using our new seminatural environment assay, we investigated social interactions pre– and post-pain in groups of five mice familiar with each other. During the nocturnal phase of the light cycle, we observed social interactions between saline-(S) and formalin-treated (F) mice. We recorded emitted and received behaviors as well as dyadic interactions. Despite targeting the neck and not the hind paw, it was possible that mice in pain would exhibit lower levels of locomotor activity, which could in turn impair their propension to encounter other mice in the SNE. F mice indeed displayed fewer zone transitions in the SNE, but this difference was not significant (t_18_ = 2.00, p = 0.061, Fig. S3A).

Formalin-treated mice were overall socially withdrawn compared to saline-treated mice. In particular, F mice emitted less social approaches and less anogenital sniffing (F_(1, 18)_ = 7.455, p = 0.014 and F_(1, 18)_ = 6.329, p = 0.022, respectively) (Fig.4A). They also received less anogenital sniffing, nose-off occurrences and were fled less often by other mice (ps < 0.038) (Fig. 4B). The number of allosniffing occurrences received by S mice peaked in the first 5-min following injection compared to that received by F mice during the same time interval (p = 0.003; interaction: F_(6, 108)_ = 2.885, p = 0.012). Interestingly, social resting behaviors (pair resting and huddling) were not affected by mouse treatment (ps > 0.396, Fig. S3B).

**Figure 4.**
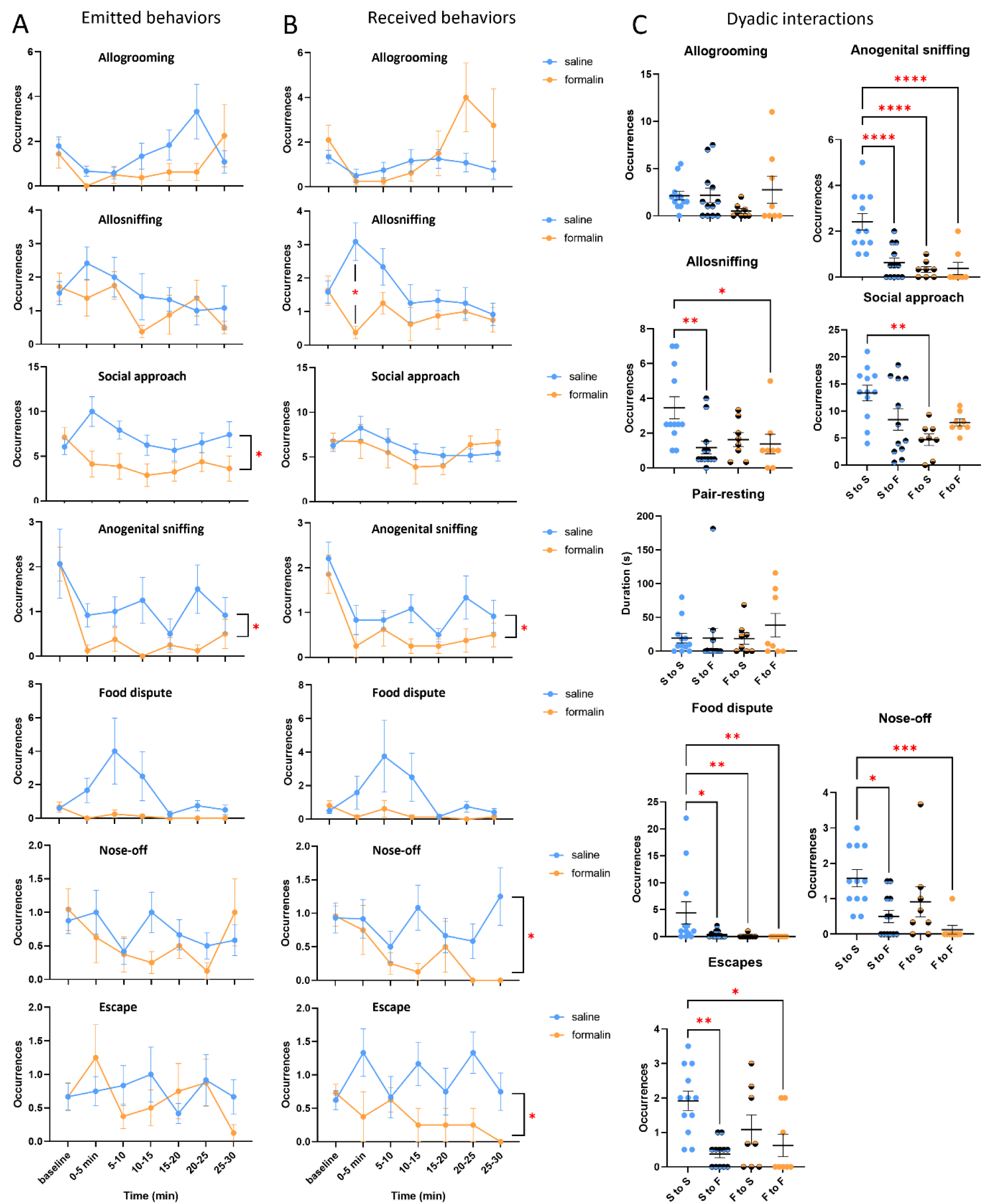
Emitted (A) and received (B) pro– and anti-social behaviors in the SNE by saline– and formalin-treated mice (blue and yellow lines, respectively). The red star on the received allosniffing plot indicates a significant difference between the treatments at the associated time interval. Other stars indicate a treatment effect. Two-way ANOVAs for repeated measured on the time interval factor, *, p < 0.05. Data are mean ± SEM. C. Dyadic interactions after formalin injection. Dyads are: saline to saline (S-S, blue), saline to formalin (S to F, half blue), formalin to saline (F to S, half yellow) and formalin to formalin mice (F to F, yellow). S to S, n=12; S to F, n=12; F to S, n=8; F to F, n=8. One-Way ANOVAs and Kruskal-Wallis rank tests. *, p < 0.05; **, p < 0.01; ***, p < 0.001; ****, p < 0.0001. Data points are dyadic interactions normalized for 1 partner by averaging the emitted behavioral occurrences.

Our observation of social withdrawal in formalin-treated mice prompted us to focus on the composition of interacting dyads, in order to decipher the directionality of social interactions. At baseline, there was no difference in social behaviors exchanged between the dyads (all ps > 0.289, Fig. S3C). However, after 2 out of 5 mice were injected with formalin, dyads including a formalin-treated mouse generally exchanged fewer social interactions. This was the case for allosniffing (F_(3, 36)_ = 4.711, p = 0.007), social approach (F_(3, 36)_ = 5.353, p = 0.004), anogenital sniffing (F_(3, 36)_ = 13.99, p < 0.001), food dispute (χ^2^ = 15.32, p = 0.002), nose-off (χ^2^ = 17.00, p = 0.001) and escape (χ^2^ = 13.74, p = 0.003). In particular, S-to-F and F-to-F dyads exchanged less allosniffing than S-to-S ones (p = 0.004 and p = 0.024, respectively). F-to-S dyads exchanged fewer social approaches than S-to-S ones (p = 0.001). All dyads including a formalin-injected mouse exchanged less anogenital sniffing than the S-to-S dyads (all ps < 0.001). This was also the case for food dispute (all ps < 0.049). Nose-off and escape behaviors were both reduced in S-to-F and F-to-F dyads compared to S-to-S ones (all ps < 0.015). Allogrooming and pair resting were the only social behaviors unaffected by dyad composition (all ps > 0.311) (Fig. 4C).

Upon our finding of social avoidance of mice in pain, we explored the link between prosocial interactions frequency and nocifensive behaviors for formalin-treated mice. We observed several negative correlations between back-scratching frequency and emitted and received prosocial behaviors during the diurnal and the nocturnal phase. Thus, we pooled diurnal and nocturnal data together. In the complete data set of 16 formalin-treated individuals thus created, back-scratching occurrences were negatively correlated to the sum of received prosocial behaviors (Spearman’s correlation, S = 1112.3, p = 0.008, rho = –0.636, sum composed of allogrooming, allosniffing, anogenital sniffing, social approach and pair-resting occurrences). The same relationship appeared with received social behavior but was not significant (S = 1010, p = 0.059, rho = –0.485) (Fig. S4). These relationships were absent at baseline (ps > 0.406).

## 3. Discussion

Mice tested in naturalistic conditions displayed less nocifensive behavior than when tested individually, consistent with our hypothesis of social buffering. However, formalin-treated mice emitted and received fewer social behaviors, attesting of pain-induced social withdrawal. Furthermore, healthy mice did not initiate social interactions with formalin-treated mice as often as with healthy controls, suggesting social avoidance of mice in pain.

Our observation that mice in pain are avoided by other mice appears contradictory to a growing body of work demonstrating empathy and potentially altruistic prosocial behaviors in rodents. However, it has repeatedly been demonstrated that observer mice exposed to a conspecific in pain show hypernociception and stress-induced analgesia [5, 10, 16, 17]. Furthermore, following approach to a conspecific in pain, mice develop place aversion [18]. Therefore, proximity with a suffering conspecific represents a significant cost for the observer, which might explain social avoidance in an environment allowing paced interactions. Accordingly, it was earlier proposed that mice approach others in pain solely to mitigate the effects of emotional contagion affecting themselves [19]. Nevertheless, findings from instrumental learning paradigms suggest that prosocial behaviors are motivated by the reduction of a conspecific’s distress [20]. In this regard, perhaps most interesting is that despite the possibility for the control mice to avoid the formalin-treated ones in the SNE, social interactions were not abolished. Though it could be argued that formalin-treated mice could not completely be avoided, thus resulting in residual social interaction, this cannot be applied to allogrooming behavior. Moreover, the duration of pair-resting, a behavior involving body contact, was comparable between formalin-treated and saline-treated animals. Those specific behaviors may carry altruistic value when displayed spontaneously in an environment allowing animals to express a wide range of their behavioral repertoire.

From our data and previously published studies, it is evident that laboratory rodents are capable of pro-social, comforting behaviors. However, in a resource-rich environment where a number of other choices are present for the pain-free mice, including exploring, eating or interacting with a healthy animal, social avoidance may be preferred. It was recently demonstrated that rats raised in complex housing conditions were less motivated to free a trapped cagemate than those raised in standard conditions [21]. This underlines the impact of physical, rather than social enrichment on prosocial behaviors. To an extent, this might even relate to human studies of altruism and compassion showing less pro-social behaviors in individuals of higher socio-economic status [22].

Notably, mice expressed fewer pain responses in the SNE than they did when isolated in a cubicle. Long-term physical enrichment has been shown to reduce pain sensitivity and promote resilience from surgically induced mechanical allodynia [23, 24], an effect potentially relying on physical activity [25]. Exploratory behaviors in the form of exploratory sniffing and nesting persisted in the SNE even in mice injected with formalin, while these behaviors were strongly diminished in the individual cubicle. Thus, high levels of exploration, providing physical activity and novelty stimulation, can have down-modulated pain behavior. Social enrichment can also alleviate pain responses via social buffering with familiar individuals [26]. Dyadic observations of a freely moving mouse and a jailed cagemate injected with acetic acid reported a negative correlation between the amount of social contact and writhing behavior [3]. However, the directionality of this relationship is unclear. It is possible that social proximity provides pain relief for the affected mice, or that healthy mice avoid affected mice proportionally to the pain levels these express. In that sense, it could be argued that the decrease in nocifensive behavior observed in the SNE compared to individually housed mice may not be due to social buffering, but rather to mice concealing their internal state to healthy conspecifics. This, however, appears unlikely as it would not only necessitate the mouse to project its internal state onto its conspecifics, but also to infer their behavioral response and how it would in turn be itself affected by this response. In addition, decreases in pain and fear behavior in the presence of a familiar conspecific have been consistently linked to a decrease in glucocorticoids plasma levels [26, 27]. It is therefore most probable that the observed reduction in nocifensive behavior in the SNE is due to social buffering.

We used a formalin pain model to investigate the link between pain levels and social behavior. When injected into the hind paw, formalin caused several changes with respect to facial expression and nocifensive behaviors, consistent with the classical two-phase pattern of this assay (see also Le Moëne and Larsson [15]). Unexpectedly, when injected into the hairy skin of the neck, this pattern was absent. This is consistent with a recent study targeting the hairy hind leg of rats with formalin, in which the first phase of nocifensive behaviors was not detected [28]. Our results showed nocifensive back-scratching peaking 10 min after injection, consistent with the beginning of formalin’s last phase. Interestingly, immobility peaked in the first 5 min. This peak might have been reinforced by the effect of anesthesia, but its absence in saline-treated animals suggests that the pain was instead expressed through freezing in the first 5 min. The mechanisms by which formaldehyde triggers nociceptive signaling and pain are diverse, and whether the observed location dependence in the dynamics of formalin-induced nocifensive behavior can be attributed to differences between glabrous and hairy skin with regards to innervation or tissue structure is unclear. Nevertheless, targeting the nape of the neck successfully elicited a pain face and pain behavior in mice, validating its use as a pain model.

## 4. Conclusion

After seven days of group-housing and in a short time-window of 60 min, we saw a clear reorganization of group and dyadic interactions following formalin injection. We established that mice housed in a socially and physically enriched seminatural environment displayed less pain behavior, likely due to social buffering in conjunction with the possibility to do more physical activity and to receive more novelty stimulation. Interestingly, saline-treated mice avoided those treated with formalin. It is possible that when given the choice, mice simply prefer to engage in more rewarding behaviors. These results finely illustrate the reciprocal influence of pain on social behavior.

## 5. Material and methods

### 2.1 Animals

Adult C57BL/6JRj mice from Janvier Laboratories were divided into two experiments: individual observation (40 mice) and group observation (40 mice). Mice were an average of 2.5 months old (10.45 ± 0.35 weeks) and weighed 21.77 ± 0.37 g. The animals lived in groups of 2-4 same-sex individuals in NexGen IVC system Mouse 500 home cages prior to the experiment, in a 20±2°C environment, under a 12L:12D light cycle. The nocturnal phase lasted from 19.00 to 7.00.

### 2.2 Apparati

The observation cubicle was a box of transparent Plexiglas of dimensions 9L x 5H x 5W, pierced with holes on the roof or the side for ventilation. The cubicle was cleaned in between each individual session.

The seminatural environment (SNE) consisted of an open area (H50 x L45 x W30 cm) communicating with a burrow area through two small openings. The burrow area was composed of a big nest box (H10 x L15 x W15 cm) linked to three small ones (H10 x L8 x W8 cm) by transparent tunnels of 3 cm diameter (GEHR, PA, USA). Both the open and burrow areas were made of opaque black Plexiglas, while the tunnels and nest boxes were made of transparent one (Interglas, Gothenburg, Sweden) (Fig. S1). Enrichment was provided in the form of nesting material, 4 wood biting sticks, 5 mats of non-woven fibers, and 2 red polycarbonate rodent tunnels. Food and water were available *ad libitum*. The burrow area was maintained in complete darkness with the help of an infrared-transmitting black lid (962 perspex IR), as suggested in Bove, Ike [29]. Both the cubicle and the SNE were submitted to the same temperature and light cycle as the home cages. In addition to the light cycle, an infrared lamp provided sufficient light for video recording.

### 2.3 Procedure

#### 2.3.1 Individual observations: Cubicle

Mice in the individual observation group were singly placed in the cubicle without any stimulation for 30 min to establish a baseline. They returned to their home cage for 5 min, allowing for feeding and drinking. They were then briefly anesthetized and injected with either saline or formalin, to be returned to the cubicle for 30 more min of observation. Anesthesia was done under isoflurane inhalation (3.5 %) and lasted no more than 60 s. Control groups received 20 μl saline (9 mg/ml) intradermally into the right hind paw (10 mice, SH group) or the nape of the neck (10 mice, SN group). Treatment groups received either 20 μl formalin (2%) into the right hind paw (10 mice, FH group) or the nape of the neck (10 mice, FN group). Each group contained 5 females and 5 males. The observations were recorded with either a monochrome Basler ace camera (a2A2590-60umPRO) or a color FLIR camera (Blackfly BFS-U3-23S3C-C) with a 1:1.8/4 mm Basler lens (C125-0418-5M) All experiments were done between 9.00 and 14.00. Mice were killed following experiment completion.

#### 2.3.2 Group observations: SNE

Groups of 5 mice (4 females + 1 male) were released in the SNE on Day 0 and left undisturbed until Day 5. On Day 5, mice were captured a first time, lightly anesthetized and all given 20 μl saline intradermally into the nape of the neck. They were then released back into the SNE. One hour later, the mice were re-captured and re-anesthetized. This time, 3 mice (2 females + 1 male) received saline again, while 2 mice received formalin (2%, 20 μl). A drop of formalin was deposited on the neck of saline-injected mice to avoid the confound of odor. The mice were then released back into the SNE and left undisturbed until day 6 when the experiment was terminated and the mice killed. The entire duration of the stay in the SNE was recorded with a monochrome Basler ace camera (a2A2590-60umPRO) and saved with the Open Broadcaster Software 27.0.1 (https://obsproject.com/). Two SNE groups were run in parallel.

Four groups of mice (n = 20) were treated during the diurnal phase of Day 5 in the SNE, in order to be compared to mice tested in the cubicle, which experiment was also conducted during the diurnal phase. In this case, mice in the SNE were observed between 9.00 and 11.00. Four more groups were treated during the nocturnal phase, when mice were most active, in order to investigate social interactions. These mice were then observed between 19.30 and 21.30.

### 2.4 Data collection

Facial expressions were assessed by measuring 7 facial parameters on mouse profiles: eye opening, ear opening, ear angle, ear position, snout position, mouth position and face inclination. These parameters have been defined and established in a previous study conducted at our lab (Le Moëne and Larsson, 2023). Briefly, from the video recordings, a lateral view picture was extracted every minute, accounting for 30 pictures per video. Facial parameters were then measured manually with IC measure (https://www.theimagingsource.com/). The side of the face that was captured on the pictures (left or right) was recorded.

Behaviors displayed in the cubicle and in the SNE were manually scored with The Observer XT 15 (Noldus). Some behaviors were displayed in both apparati, but some could only be expressed in the SNE. The ethogram used is visible in Table 1. In the cubicle, behaviors were scored for the entire 30-min of baseline and post-formalin injection. In the SNE, mouse behavior was scored for 30 min following the first injection, and for 30 more minutes following the second injection. Locomotor behavior was measured by the frequency of zone transitions, after dividing the SNE into four zones of equal surface in the open area, and four more zones in the burrow area corresponding to the main nest box and the three smaller ones with their associated tunnels.

**Table 1.** Ethogram used in the cubicle and the seminatural environment. f = behavior scored in frequency; d = behavior scored in duration. SNE = Behaviors only scored in the SNE.

| Category | Behavior pattern | Description |
| --- | --- | --- |
| Exploratory behaviors | Exploratory sniffing; f | Sniffs the environment with all four paws on the floor. |
|  | Rearing; f; SNE | Sniffs the air while standing on hind legs. |
|  | Stretch-attend posture; f; SNE | Stops on-going behavior to adopt a vertically stretched posture, ears pointing to the object of attention. |
| Maintenance and resting behaviors | Self-grooming; f | Grooms its own body with front paws. |
|  | Nesting; f | Transports or moves nest material or excrements. |
|  | Eating; f; SNE | Self-explanatory. |
|  | Zap; f; SNE | Sudden contraction of the muscles resulting in an unprompted startle or a rapid change in locomotion direction. |
|  | Immobility; d | Stays immobilized, includes sleeping alone or next to other animals. |
|  | Pair-resting; f; d; SNE | Rests in body contact with one other mouse. |
|  | Huddling; d; SNE | Rest in body contact with two mice or more. |
| Prosocial behaviors | Approach; f; SNE | Approaches and sniffs another animal who does not reciprocate the sniffing. |
|  | Allosniffing; f; SNE | Sniffs another mouse that reciprocates the sniffing. |
|  | Allogrooming; f; SNE | Grooms another mouse. |
|  | Anogenital sniffing; f; SNE | Sniffs the anogenital region of another mouse. |
| Agonistic behaviors | Nose-off; f; SNE | Faces another mouse either standing on four legs or rearing; includes boxing and teeth showing. |
|  | Escape; f; SNE | Escapes from an interaction or avoids social interaction by moving head aside or walking/running away. |
|  | Food dispute; f; SNE | Approaches another individual eating and pulls the food pellet to itself. |

### 2.5 Data preparation and statistics

#### 2.5.1 Facial expressions

First, we divided the 30-min of baseline and post-injection phases into six time intervals of 5 min each. The value at a time interval corresponds to the average of the 5 frames captured in this 5-min time intervals. Upon returning to the cubicle following injection, some of the mice injected with saline fell asleep. Associated frames were thus excluded to avoid confounding the facial expressions. Then, to create facial response profiles, we calculated for each facial parameter, for each experimental group (SH, FH, SN and FN) its proportional change from baseline with the formula: (value at stimulus – value at baseline)/value at baseline. Differences from baseline were analyzed by one-sample *t*-tests.

#### 2.5.2 Behaviors

We analyzed behavioral changes in mice undergoing injection into the paw or the neck in the observation cubicle or the SNE. Data are presented as the sum of occurrences in 5-min intervals across the baseline and post-injection. Then, we compared the effect of the apparatus on behavioral dynamics. A single average baseline value per individual was computed from the average of all 6 baseline time intervals. Analyses were conducted separately for saline-treated mice (SN-cubicle and SN-SNE) and for formalin-treated mice (FN-cubicle and FN-SNE).

In the SNE, since each mouse group was composed of 3 saline-treated (S) and 2 formalin-treated (F) individuals, each S individual could interact with 2 F individuals while each F individual could interact with 3 individuals. Therefore, dyadic interactions were normalized for 1 partner by averaging the emitted behavioral occurrences.

Statistical analyses are detailed in figure captions. Plots and statistics were obtained with GraphPad Prism 9, RStudio and R 4.1.3 (core, tidyverse, and scales packages).

## Acknowledgements

Knut and Alice Wallenberg Foundation, project no. 2019.0047.

## Author contributions

OLM and ML designed the experiment, OLM performed the experiment, OLM and ML analyzed the data, OLM and ML wrote the paper.

## Data and materials availability

All data and codes are available on https://github.com/olm976/pain_social_behavior

## Supplementary Materials

**Figure S1.**
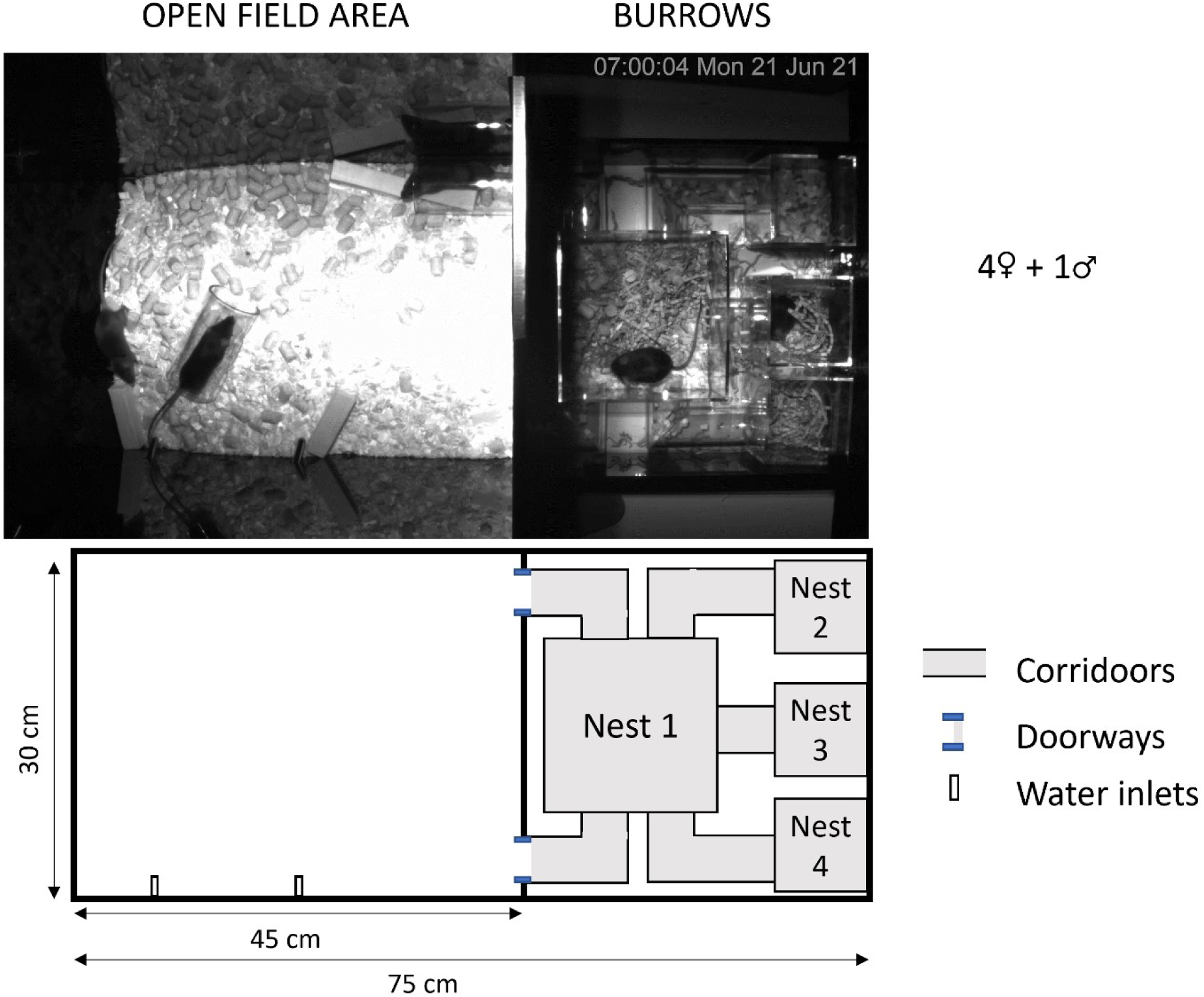
Picture of the seminatural environment (SNE) (top). Scheme of the SNE (bottom). Five mice (4 females and 1 male) are placed in the SNE for 7 days (day 0 to 6) with food and water available *ad libitum*.

**Figure S2.**
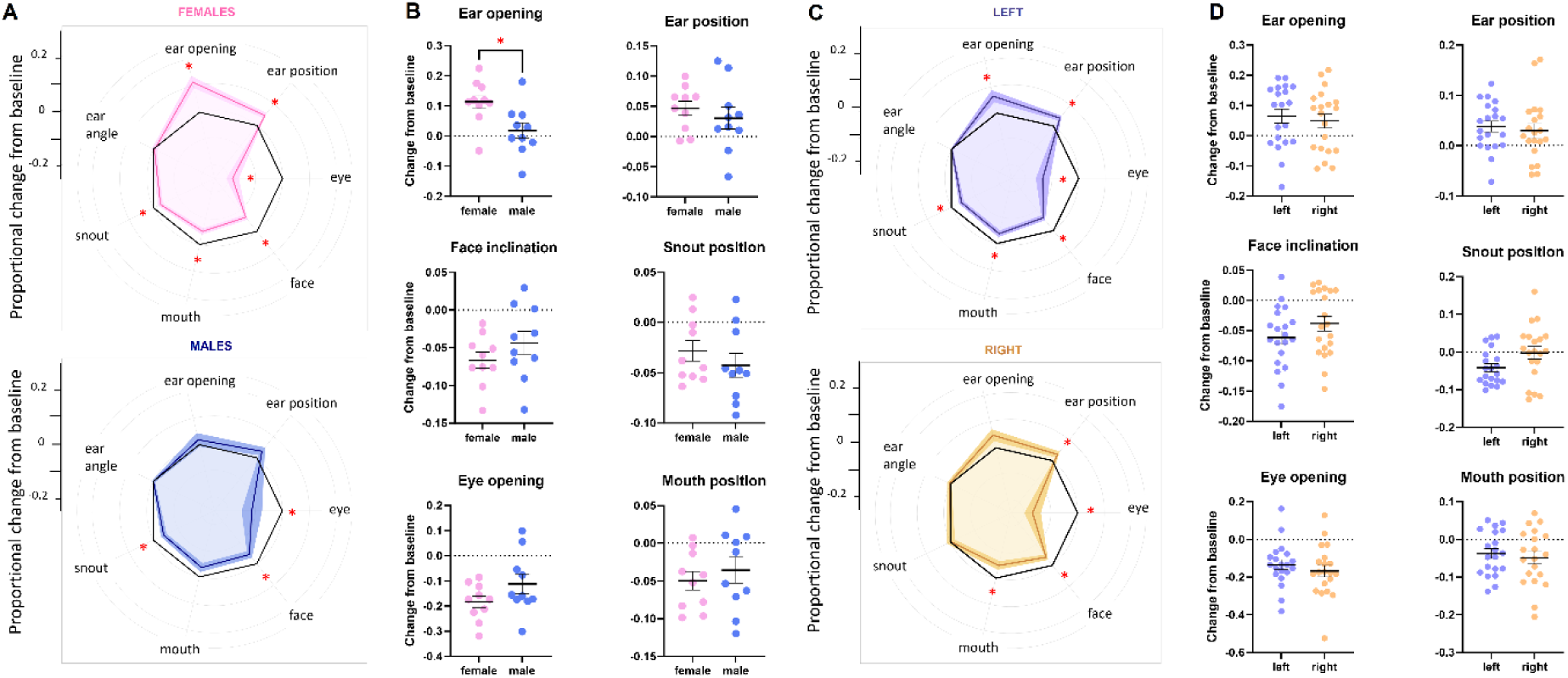
Facial expressions elicited by formalin injection (neck and hind paw data pooled, n = 10 females + 10 males). A. Female (pink) and male (blue) response profile to formalin (30 min observation). B. Sex difference in facial parameters’ change from baseline. C. Responses profiles to formalin produced by pictures from the left (purple) and right (yellow) side of mouse faces. D. Side difference in facial parameters’ change from baseline. Data are mean ± SEM and individual data points. Response profiles (A-C): one-sample *t*-test; Scatter plots (B-D): *t*-test. *, p < 0.05.

**Figure S3.**
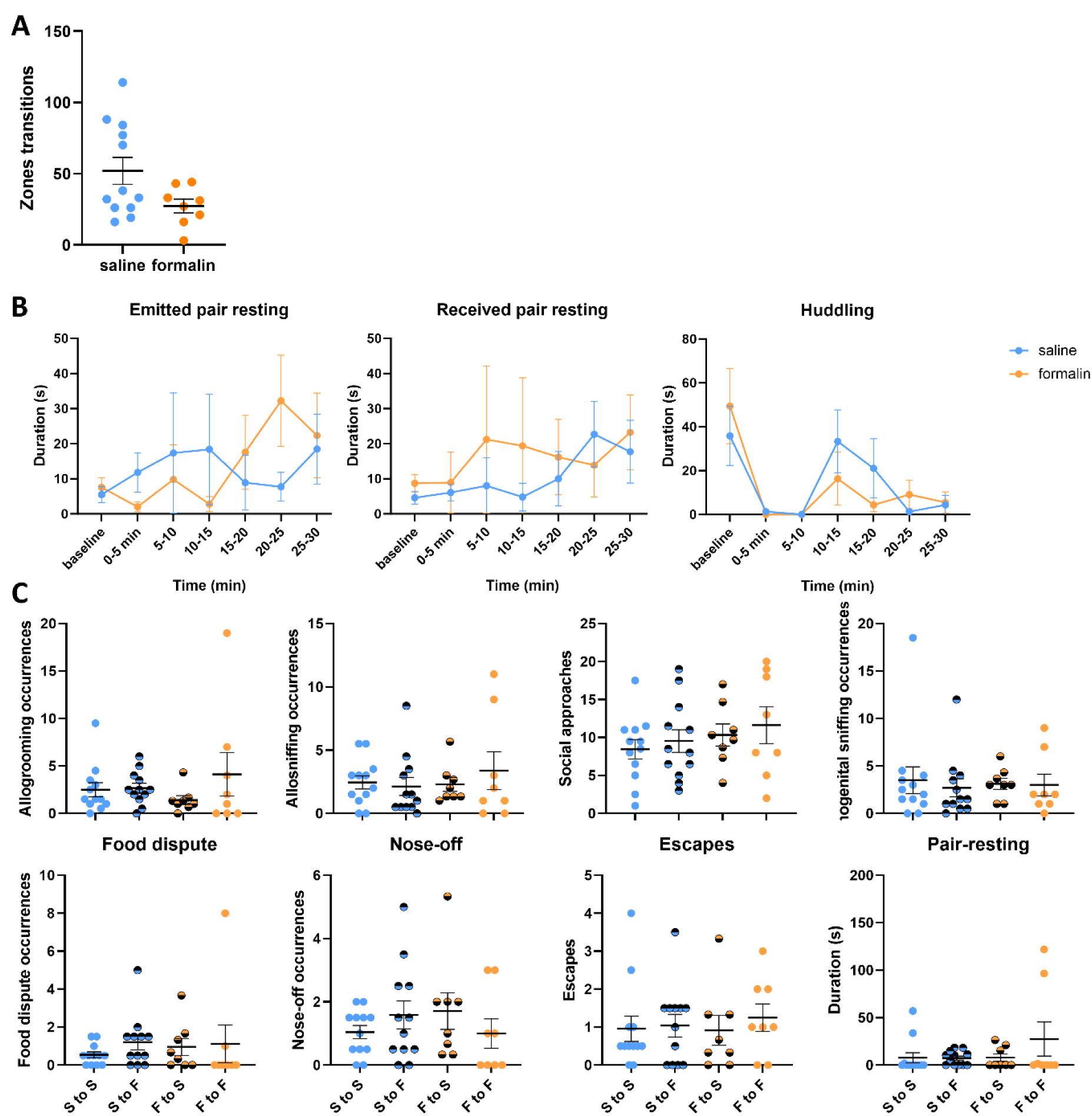
**A**. Zone transitions in the SNE, *t*-test **B**. Emitted and received resting behaviors. Two-way ANOVAs for repeated measured on the time interval factor, *, p < 0.05. Data are mean ± SEM. **C**. Dyadic interactions at baseline from saline to saline (S-S, blue, n=12), saline to formalin (S to F, half blue, n=12), formalin to saline (F to S, half yellow, n=8) and formalin to formalin (F to F, yellow, n=8) mice. One-Way ANOVAs and Kruskal-Wallis rank tests, all ps > 0.289.

**Figure S4.**
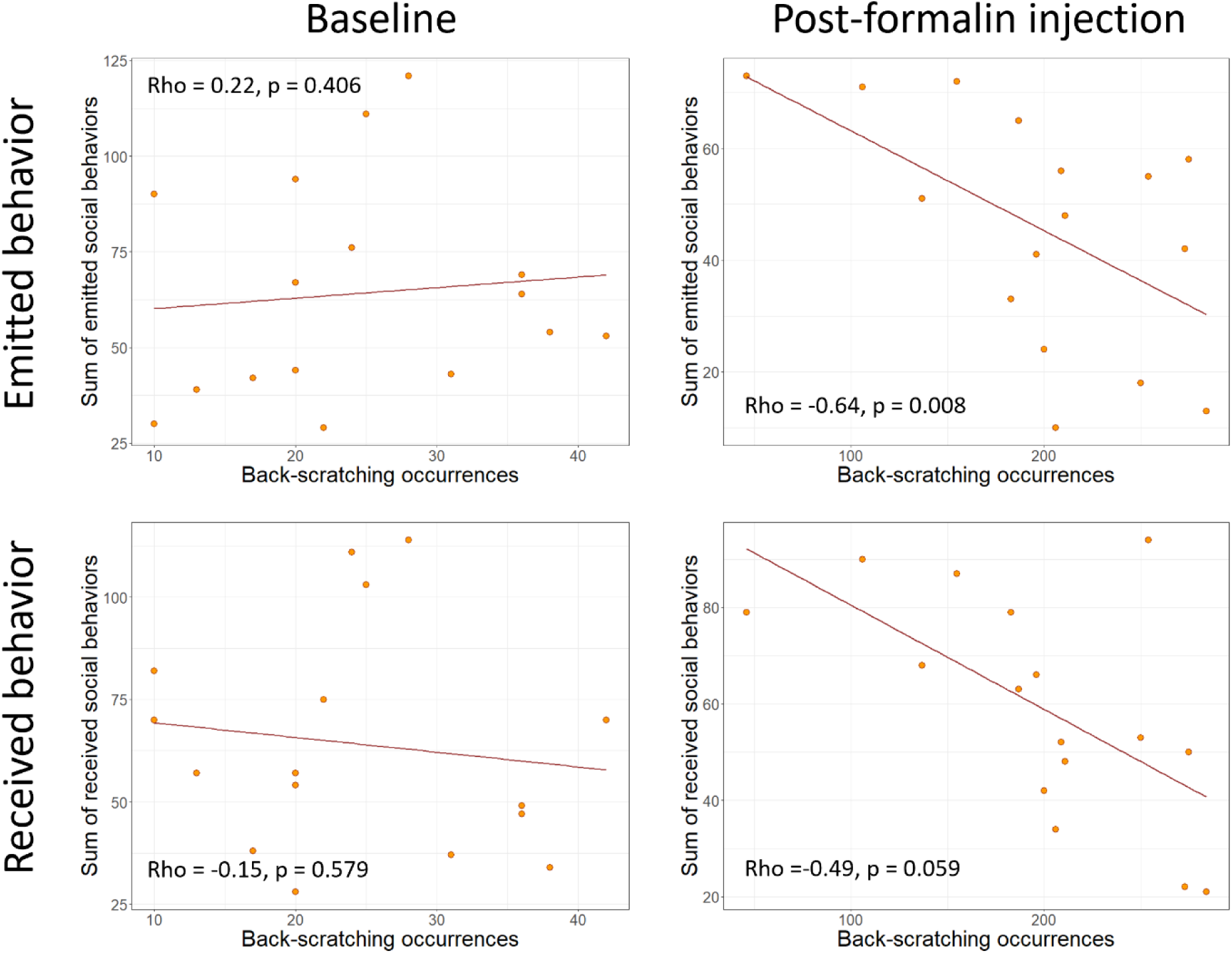
Spearman’s correlations between occurrences of back-scratching and the sum of emitted (top row) or received (bottom row) social behavior occurrences at baseline (left) or following formalin injection (right). Included social behaviors are allogrooming, allosniffing, anogenital sniffing, social approach and pair-resting. N=16 formalin-treated mice (8 during the diurnal and 8 during the nocturnal phase of the light cycle).

## References

1. Mogil, J.S., Social modulation of and by pain in humans and rodents. Pain, 2015. 156 **Suppl 1**: p. S35–S41.

2. Kiyokawa, Y. and M.B. Hennessy, Comparative studies of social buffering: A consideration of approaches, terminology, and pitfalls. Neurosci Biobehav Rev, 2018. 86: p. 131–141.

3. Langford, D.J., et al., Social approach to pain in laboratory mice. Soc Neurosci, 2010. 5(2): p. 163–70.

4. Hostinar, C.E., R.M. Sullivan, and M.R. Gunnar, Psychobiological mechanisms underlying the social buffering of the hypothalamic-pituitary-adrenocortical axis: a review of animal models and human studies across development. Psychol Bull, 2014. 140(1): p. 256–82.

5. Baptista-de-Souza, D., et al., Behavioral, hormonal, and neural alterations induced by social contagion for pain in mice. Neuropharmacology, 2022. 203: p. 108878.

6. Brunswik, E., Representative design and probabilistic theory in a functional psychology. Psychol Rev, 1955. 62(3): p. 193–217.

7. Huzard, D., et al., The impact of C-tactile low-threshold mechanoreceptors on affective touch and social interactions in mice. Sci Adv, 2022. 8(26): p. eabo7566.

8. Blanchard, D.C., et al., Visible burrow system as a model of chronic social stress: Behavioral and neuroendocrine correlates. Psychoneuroendocrinology, 1995. 20(2): p. 117–134.

9. Le Moëne, O. and A. Ågmo, Behavioral responses to emotional challenges in female rats living in a seminatural environment: The role of estrogen receptors. Horm Behav, 2018. 106: p. 162–177.

10. Baptista-de-Souza, D., et al., Mice undergoing neuropathic pain induce anxiogenic-like effects and hypernociception in cagemates. Behav Pharmacol, 2015. 26(7 Spec No): p. 664–72.

11. Langford, D.J., et al., Social Modulation of Pain as Evidence for Empathy in Mice. Science, 2006. 312(5782): p. 1967–1970.

12. de Waal, F.B.M. and S.D. Preston, Mammalian empathy: behavioural manifestations and neural basis. Nat Rev Neurosci, 2017. 18(8): p. 498–509.

13. Russell, Y.I., Allogrooming, in Encyclopedia of Animal Cognition and Behavior, J. Vonk and T. Shackelford, Editors. 2017, Springer International Publishing: Cham. p. 1–4.

14. Moser, R., M. Cords, and H. Kummer, Social influences on grooming site preferences among captive long-tailed macaques. International Journal of Primatology, 1991. 12(3): p. 217–230.

15. Le Moëne, O. and M. Larsson, *A New Tool for Quantifying Mouse Facial Expressions*. eneuro, 2023. 10(2): p. ENEURO.0349-22.2022.

16. Lu, Y.F., et al., Social interaction with a cagemate in pain increases allogrooming and induces pain hypersensitivity in the observer rats. Neurosci Lett, 2018. 662: p. 385–388.

17. Martin, L.J., et al., Reducing social stress elicits emotional contagion of pain in mouse and human strangers. Curr Biol, 2015. 25(3): p. 326–332.

18. Watanabe, S., Distress of mice induces approach behavior but has an aversive property for conspecifics. Behavioural Processes, 2012. 90(2): p. 167–173.

19. Rice, G.E. and P. Gainer, “Altruism” in the albino rat. J Comp Physiol Psychol, 1962. 55: p. 123–5.

20. Keysers, C., et al., Emotional contagion and prosocial behavior in rodents. Trends Cogn Sci, 2022. 26(8): p. 688–706.

21. Kalamari, A., et al., Complex Housing, but Not Maternal Deprivation Affects Motivation to Liberate a Trapped Cage-Mate in an Operant Rat Task. Front Behav Neurosci, 2021. 15: p. 698501.

22. Piff, P.K. and J.P. Moskowitz, The class–compassion gap: how socioeconomic factors influence compassion, in The Oxford handbook of compassion science, E.M. Seppälä, et al., Editors. 2017, Oxford University Press. p. 317–330.

23. Falkowska, M., et al., Environmental enrichment promotes resilience to neuropathic pain-induced depression and correlates with decreased excitability of the anterior cingulate cortex. Front Behav Neurosci, 2023. 17: p. 1139205.

24. Kimura, L.F., V.G.M. Mattaraia, and G. Picolo, Distinct environmental enrichment protocols reduce anxiety but differentially modulate pain sensitivity in rats. Behav Brain Res, 2019. 364: p. 442–446.

25. Gabriel, A.F., et al., Enriched environment and the recovery from inflammatory pain: Social versus physical aspects and their interaction. Behav Brain Res, 2010. 208(1): p. 90–5.

26. Denommé, M.R. and G.J. Mason, Social Buffering as a Tool for Improving Rodent Welfare. Journal of the American Association for Laboratory Animal Science, 2022. 61(1): p. 5–14.

27. Kiyokawa, Y., K. Kawai, and Y. Takeuchi, The benefits of social buffering are maintained regardless of the stress level of the subject rat and enhanced by more conspecifics. Physiol Behav, 2018. 194: p. 177–183.

28. Erami, E., et al., Characterization of Nociceptive Behaviors Induced by Formalin in the Glabrous and Hairy Skin of Rats. Basic Clin Neurosci, 2017. 8(1): p. 37–42.

29. Bove, M., et al., The Visible Burrow System: A behavioral paradigm to assess sociability and social withdrawal in BTBR and C57BL/6J mice strains. Behav Brain Res, 2018. 344: p. 9–19.

